# Acinetobase: the comprehensive database and repository of *Acinetobacter* strains

**DOI:** 10.1101/2022.02.27.482141

**Authors:** Adam Valcek, James Collier, Alexander Botzki, Charles Van der Henst

## Abstract

**Background:** *Acinetobacter baumannii* is one of the most problematic nosocomial pathogens that can efficiently thrive within hospital settings, mainly due to resistances towards antibiotics, desiccation, disinfectants, human serum and oxidative stress. Recently, increased resistance against last resort antibiotics earns this bacterium the highest priority concern classified by the CDC and the WHO. An obvious hallmark of this bacterium is the high heterogeneity observed amongst *A. baumannii* isolates, with a limited core genome. This feature complexifies the study of *A. baumannii* bacteria as an entity, subsequently reflected in a diversity of phenotypes of not only antimicrobial and environmental resistance but also virulence.

**Main body:** High degree of genome plasticity, along with the use of a limited subset of established reference strains, can lead to strain-specific observations, decreasing the global understanding of this pathogenic agent. Phenotypic variability of *A. baumannii* strains is easily observable such as with the macrocolony morphologies, *in vitro* and *in vivo* virulence, natural competence level, production of different capsular polysaccharide structures and cellular densities. Some strains encode an extensive amount of virulence factors while other, including the reference strains, lack several key ones. The lack/excess of genes or specific physiological processes might interfere with *in vivo* and *in vitro* experiments, thus providing limited impact on the global understanding of *Acinetobacter* bacteria as a whole.

**Conclusion:** As an answer to the high heterogeneity amongst *A. baumannii* strains, we propose a first comprehensive database that includes the bacterial strains and the associated phenotypic and genetic data. This new repository, freely accessible to the entire scientific community, allow selecting the best bacterial isolate(s) related to any biological question, using an efficient and fast exchange platform.

## Background

*Acinetobacter baumannii* is an opportunistic nosocomial pathogen thriving in hospital environment and endangering especially critically ill patients (Whiteway *et al*., 2022). This Gram-negative bacterium is part of the most problematic ESKAPE group of human pathogens, against which therapeutic approaches become limited (Pendleton *et al*., 2014). Increased isolations of antibiotic-resistant strains, especially to last resort antibiotics, earns this bacterium the consideration as “urgent threat to public health” by the Centers for Disease Control and Prevention (CDC) (CDC. Antibiotic Resistance Threats in the United States and U.S. Department of Health and Human Services) and as “a bacterium for which research and development of new antibiotics is critically needed” by the World Health Organization (WHO) (Tacconelli *et al*., 2018). An obvious hallmark of *A. baumannii* bacteria is the high heterogeneity observed amongst the isolates [5]. *A. baumannii* bacteria have a dynamic genome, with an estimated conserved core genome of only 16.5%, while 25% of the genome is unique to each strain, meaning that there is no counterpart in any other *A. baumannii* genome (Imperi *et al*., 2011). Our recent study (Philippe *et al*., 2022) on 47 strains of *A. baumannii* shows even lower core genome of 2009 genes from the pangenome of 13611 genes, accounting for 14.76%. *A. baumannii* isolates are impressively variable, at both genetic and phenotypic levels. For example, the capsular polysaccharide, which impacts antibiotic and environmental resistances, host response and virulence, is encoded by at least 137 various capsule locus types, while the outer core of lipooligosaccharide by at least 13 locus types (Wyres *et al*., 2020; Kenyon and Hall, 2021). This genetic variability translates into phenotypic heterogeneity, observable using electron microscopy where the thickness and rigidity of the capsule differs, along with different cellular densities (Whiteway *et al*., 2021). The established reference strains such as ATCC19606, ATCC17978, DSM30011 or AB5075 are widely used type strains from which significant and validated observations were and are still generated (Roussin *et al*., 2019; Bravo *et al*., 2016; Jacobs *et al*., 2014; de Silva *et al*., 2017). All together, these reference strains greatly contributed to the state of the art of the current and growing *A. baumannii* field. The reference strains AB5075, ATCC17978 and ATCC19606 are multidrug resistant (MDR) strains of clinical origin [17, 18] while DSM30011 is a non-MDR environmental strain obtained from plant microbiota [19]. The observations gained about these reference strains shed a strong base of the knowledge about *A. baumannii*. However, in the other hand, type strains also showed the limitations of only using these strains when current clinical isolates of *A. baumannii* show extreme genetic and phenotypic variabilities (Philippe *et al*., 2022). In order to fill this modern gap of knowledge, grasp the complexity of *A. baumannii* and widen the possibilities of the community to use and efficiently share genotypically and phenotypically characterized strains, we propose the database and strain repository Acinetobase as a new community tool (Fig. 1). This new data and strains sharing platform is especially important in the context of novel therapeutic discoveries, for which conserved targets and antimicrobial mode of actions amongst the highest proportion of problematic and clinically relevant *A. baumannii* isolates is required.

**Figure 1:**
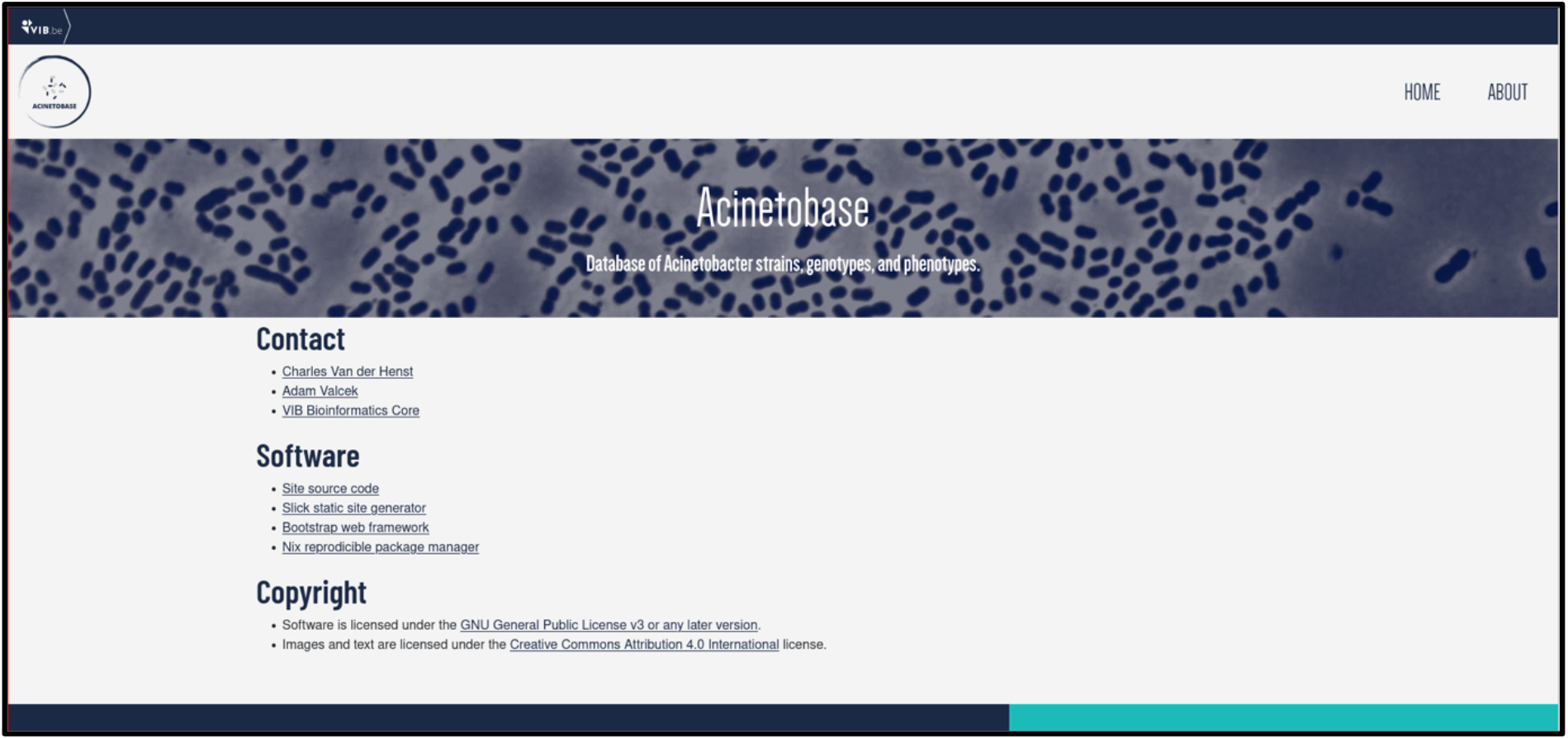
The view of the “About” page of Acinetobase.

## Construction and content

This first version of Acinetobase consists of 43 clinical isolates and four reference strains (AB5075-VUB, ATCC19606-VUB, ATCC17978-VUB and DSM30011-VUB) of *A. baumannii*. We generated whole genome sequences and *de novo* assembled the complete chromosomal sequences with high quality of all these strains, rendering them publicly available via GenBank accession number. In addition, the Acinetobase includes a variety of phenotypical characteristics associated with a strain:

i. Quantification of the capsular polysaccharide using gradient density method for each strain [12]
ii. Type of macrocolony produced for each strain (designated with letter A to E) with a corresponding photograph
iii. Virulence in *Galleria mellonella* infection model for 9 strains [12]
iv. Transmission electron microscopy (TEM) photograph depicting the amount and density of the capsule for 11 strains

The database also describes basic genotypical characteristics of each strain:

i. Multilocus sequence type (ST) according to Pasteur [20] and Oxford [21, 22] typing scheme
ii. Capsule locus type (KL) (Wick *et al*., 2018; Wyres *et al*., 2020)
iii. Outer core lipooligosaccharide locus type (OCL) (Wick *et al*., 2018; Wyres *et al*., 2020)

The above mentioned genotypical and phenotypical traits can be used to search the database and filter the strains for specific criteria. This way users can efficiently find the strain of desired characteristics and request it. The bacterial strains of *A. baumannii* were deposited and are available in lyophilized form as per order from the Belgian Coordinated Collection of Microorganisms/Lab of Microbiology (BCCM/LMG) (https://bccm.belspo.be/). The example of an entry of AB5075-VUB is depicted in Fig. 2.

**Figure 2:**
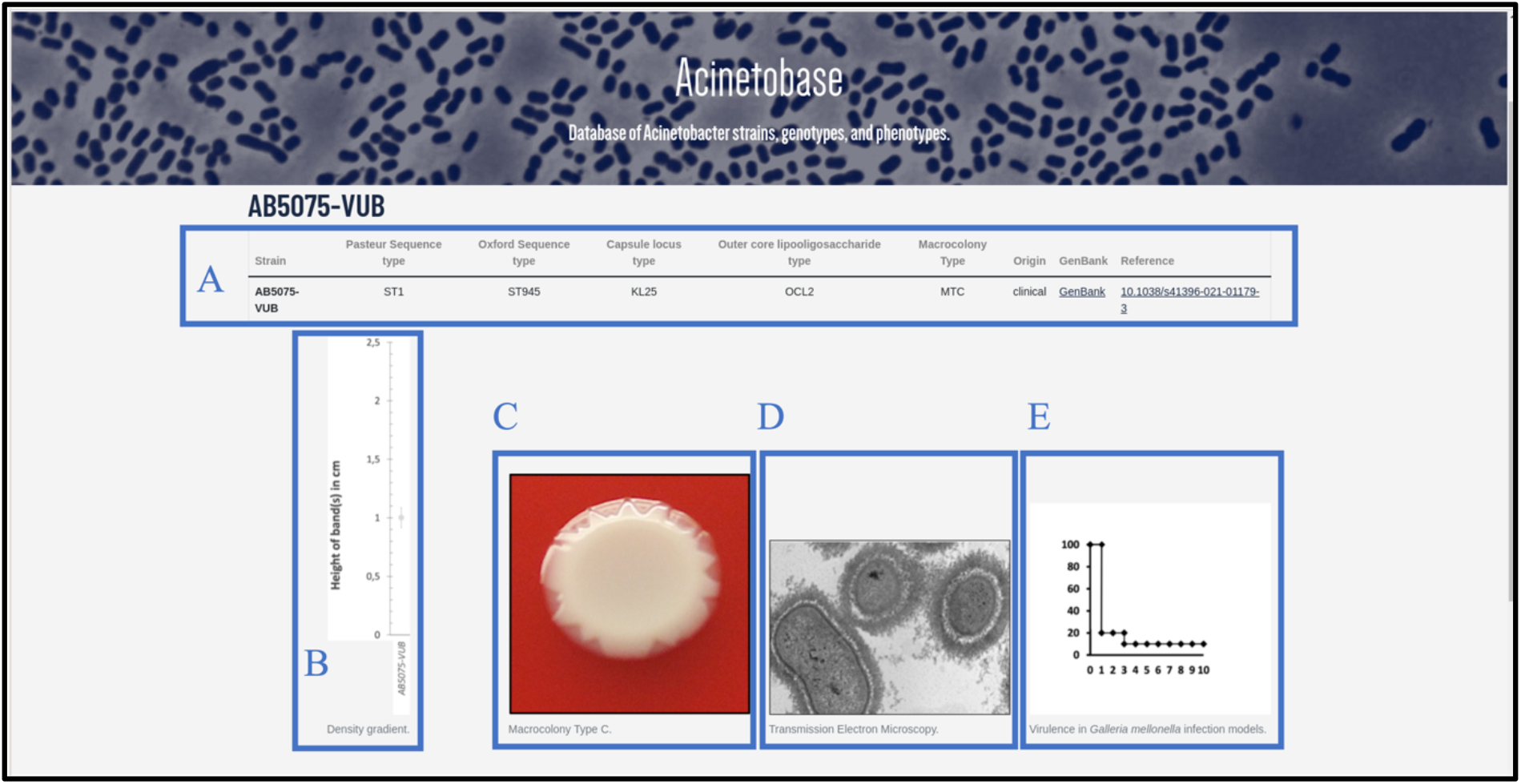
An entry of AB5075-VUB depicting **A**, description of the genotype (KL, ST^Ox^/^Pas^, OCL) with a hyperlink for the complete chromosomal sequence and a reference publication; **B**, quantification of the capsule; **C**, macrocolony type; **D**, TEM micrograph of the capsule and **E**, virulence in *Galleria mellonella* infection model.

## Utility and discussion

### Existing databases

The decreasing costs of the whole-genome sequencing allowed genomic databases, filled with high-quality data, to thrive. There are well-established databases such as EnteroBase (Zhou *et al*., 2020), which does not include data on *Acinetobacter*. In contrast, existing databases which provide whole-genome sequencing data on *A. baumannii*, such as BIGSdb (Jolley *et al*., 2018), includes metadata and genotype, however, BIGSdb does not include phenotypic data unrelated to the genotype nor the strains themselves. Moreover, centralization of the laboratory providing strains with the genotype and phenotype characteristics will help to decrease the chance of working with subcultured variants with altered genotype and phenotype, a pitfall that becomes more and more apparent in the *A. baumannii* field (Whiteway *et al*., 2021; Artuso *et al*., 2022; Wijers *et al*., 2021).

### Browsing

The users can browse the “Collection” page for the strains (Fig. 3) and pick according to their genomic description. By clicking on the name of the selected strain the full entry with genomic and phenotypic details appears (Fig. 2).

**Figure 3:**
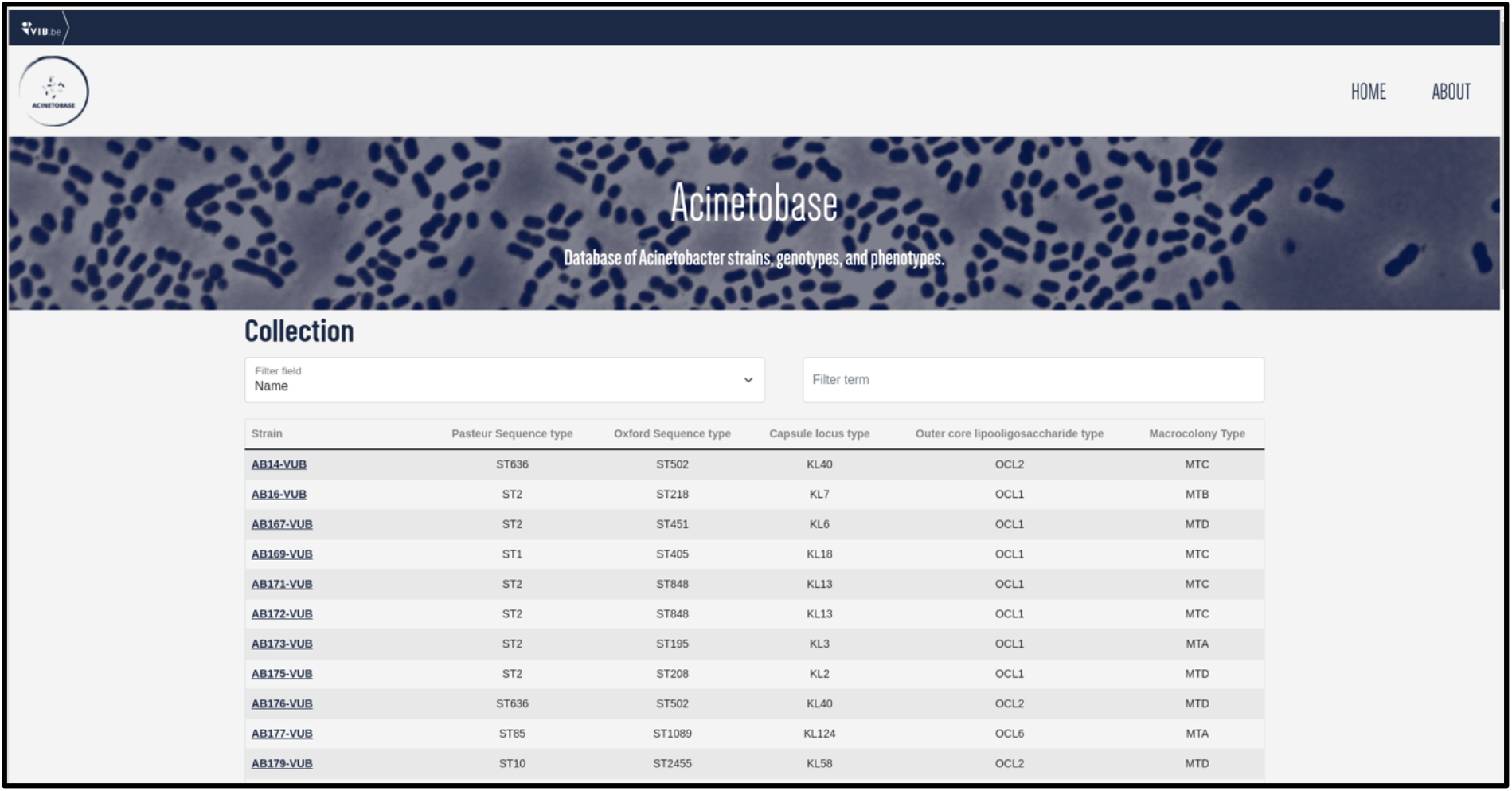
The “Collection” page depicting the name, Pasteur and Oxford sequence type, KL, OCL and macrocolony types of each isolate in the collection.

### Searching

The users can filter the entries (Fig. 4) according to each genetic and phenotypic-based criterium depicted in the “Collection” page. This feature helps to select specific strains of *A. baumannii* according to criteria demanded by the user, appropriated to their biological question.

**Figure 4:**
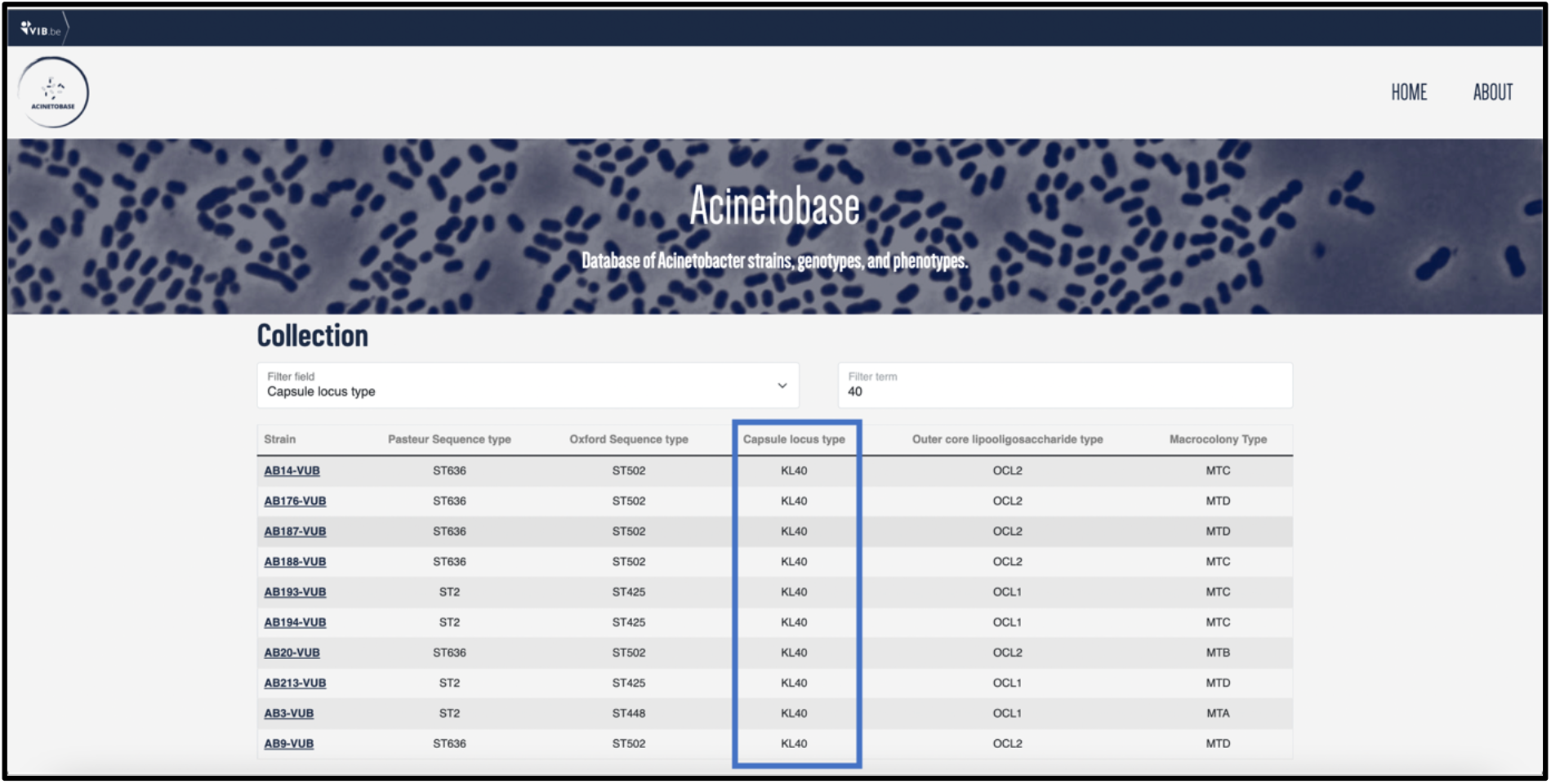
An example of filtering the entries based on the Capsular locus (KL) type 40 (in the frame).

### Download

To avoid unnecessary redundancy, downloading the complete chromosomal sequences and the whole-genome assemblies is facilitated via GenBank (https://www.ncbi.nlm.nih.gov/). The source information on the strains can be found in the corresponding research articles found in the “Reference” column using hyperlinks (using Digital Object Identifier - DOI). In the corresponding peer-reviewed study, authors of the study provide the protocol on how the strains were obtained, their origin, original figures, and the protocols for assessing the phenotypical traits. The authors of the study will be directly responsible for the information and data they provided and submitted to Acinetobase.

### Future perspectives

Acinetobase has a great potential to influence the field of microbiology, especially the *A. baumannii* community, but also other communities which work with similar pathogens and similar biological questions, might be inspired. Further implementation potential of Acinetobase resides in new strains acquisition and the accumulation of additional experimental analyses and data. This will provide valuable information to researchers whether one (or a subset of available strains) is an appropriate isolate to work with, depending of the biological question aimed. The proposed project is so far aimed to *Acinetobacter baumannii*, yet not limited to a single species of *Acinetobacter*. With an increasing clinical relevance of other *Acinetobacter* species such as *A. pittii, A. nosocomialis, A. seifertti* and *A. dijkshoorniae* (Vijayakumar *et al*., 2019), the project of Acinetobase is open to all *Acinetobacter* species. Acinetobase will be open to submission of the phenotypic and genotypic data including strains from other laboratories and research groups. This initiative will, over time, result in a growing database and a platform for sharing knowledge and *Acinetobacter* spp. strains and might include further molecular tools such as fluorescently labeled strains or characterized deletion mutant strains.

## Conclusion

In conclusion, Acinetobase is the first database dedicated to *Acinetobacter* genotypes and associated phenotypes that also serves as a repository of characterized strains. Acinetobase will not only be the proof-of-concept of high heterogeneity among *A. baumannii* strains but will transform this pitfall into a new strength, benefiting the entire community by helping to select an appropriate strain related to specific conditions, experiments, and biological questions. The subculturing steps of the current strain sharing strategy will therefore decrease, helping to keep the genetic background of these dynamic bacteria as much as possible.

## Declarations

### Ethics approval and consent to participate

Not applicable.

### Consent for publication

Not applicable.

### Availability of data and materials

The datasets generated and analyzed during the current study are linked from the Acinetobase website, http://acinetobase.vib.be/.

### Authors’ contributions

AV and CVDH wrote up the manuscript. CVDH conceptualized the Acinetobase. JC wrote and designed the website, data storage, and update mechanisms. AB assisted with design of the website, is the project sponsor and reviewed the manuscript.

## Acknowledgements

We would like to thank to Belgian Coordinated Collections of Microorganisms of Ghent University, Faculty of Sciences, Belgium and the VIB Bioinformatics Core for their support in developing the website, especially to Christof De Bo who helped with graphics. We also would like to thank the National Reference Laboratory for Monitoring of Antimicrobial Resistance in Gram-negative Bacteria, CHU Mont-Godinne, Université Catholique de Louvain (UCL), Yvoir, Belgium for sharing the modern clinical isolates of *A. baumannii*.

